# CLARA: A web portal for interactive exploration of the cardiovascular cellular landscape in health and disease

**DOI:** 10.1101/2021.07.18.452862

**Authors:** Malathi S.I. Dona, Ian Hsu, Thushara S. Rathnayake, Gabriella E. Farrugia, Taylah L. Gaynor, Malvika Kharbanda, Daniel A. Skelly, Alexander R. Pinto

**Affiliations:** Baker Heart and Diabetes Research Institute, Melbourne, Victoria, Australia; The Jackson Laboratory, Bar Harbor, ME, USA; Centre for Cardiovascular Biology and Disease Research, La Trobe University, Melbourne Victoria, Australia

## Abstract

Mammalian cardiovascular tissues are comprised of complex and diverse collections of cells. Recent advances in single-cell profiling technologies have accelerated our understanding of tissue cellularity and the molecular networks that orchestrate cardiovascular development, maintain homeostasis, and are disrupted in pathological states. Despite the rapid development and application of these technologies, many cardiac single-cell functional genomics datasets remain inaccessible for most cardiovascular biologists. Access to custom visual representations of the data, including querying changes in cellular phenotypes and interactions in diverse contexts, remains unavailable in publicly accessible data portals. Visualizing data is also challenging for scientists without expertise in processing single-cell genomic data. Here we present CLARA—CardiovascuLAR Atlas—a web portal facilitating exploration of the cardiovascular cellular landscape. Using mouse and human single-cell transcriptomic datasets, CLARA enables scientists unfamiliar with single-cell-omic data analysis approaches to examine gene expression patterns and the cell population dynamics of cardiac cells in a range of contexts. The web-application also enables investigation of intercellular interactions that form the cardiac cellular niche. CLARA is designed for ease-of-use and we anticipate that the portal will aid deeper exploration of cardiovascular cellular landscapes in the context of development, homeostasis and disease. CLARA is freely available at https://clara.baker.edu.au.

## INTRODUCTION

The heart and other cardiovascular tissues are formed by a diverse array of cell types. In the heart, these include myocytes—which form the vast majority of the heart’s volume—and non-myocytes, which outnumber myocytes. The application of single-cell flow cytometry and high throughput sequencing methodologies have transformed our understanding of cardiovascular tissue cellularity and, in particular, the heterogeneity of non-myocytes of the heart [1, 2]. Single-cell transcriptomics enables joint examination of cellular heterogeneity and gene expression patterns, in addition to revealing shifts in these parameters in context of development or tissue stress. Recently, multiple single-cell transcriptomic studies have been published—in human and non-human contexts—that have examined the cellular landscape of the heart [2–6] and aorta [7, 8]. Collectively, these studies have shown that cell types respond in an orchestrated manner to a range of physiological stressors, and that the cardiovascular tissues are ecosystems of interdependent cell types. While these studies have provided new and valuable insights into the community of cells that form these tissues, much of the data exploration and analyses presented in individually published studies have been focused on specific or narrowly defined questions confined by the scope of each study. Thus, there is a need for ready accessibility of these datasets in order to broaden their value for application to disparate research fields.

Motivated by the need for a user-friendly and accessible resource for exploration of cardiac single-cell genomic datasets, we developed CLARA (CardiovascuLAR Atlas), a web portal for exploring the cardiac cellular landscape. The CLARA portal provides the capacity to explore gene expression within individual cell populations and interactions between populations. Specific features include visualization and retrieval of: (i) expression of any gene of interest, including changes in context of physiological stress such as fibrosis and myocardial infarction; (ii) top-ranked genes that define cell populations; (iii) top-ranked genes that change within individual cell populations in the context of physiological stress; and (iv) inter-cellular ligand-receptor interactions. It is anticipated that CLARA will provide a customized and user-specific experience that will aid exploration of the cellular landscapes of tissues of interest to cardiovascular researchers.

## RESULTS

To enable analysis of the cardiovascular cellular landscape, we prepared datasets from six cardiac and aortic single-cell transcriptomic datasets published in eight independent studies. These include four mouse heart datasets (Table 1) [2–5], one human heart dataset [6], one mouse aorta [8] and one human aorta [7]. All datasets allow examination of the uninjured heart or aorta. In addition, the mouse datasets permit consideration of changes in cellularity and gene expression in a variety of pathological states, including myocardial infarction [3, 5], angiotensin II (AngII)-induced cardiac fibrosis [4], and elastase-induced aortic aneurism [8]. The mouse datasets were developed using whole intact cells, with one of them including nuclei of cardiomyocytes [4], while the human datasets were generated from cell nuclei [6, 7]. We also provide a computationally integrated dataset consisting of four mouse heart datasets merged together [2–5], which serves as a unified portrait of changes to the heart across a variety of pathological conditions.

**Table 1.** Summary of datasets used in CLARA.

|  | Skelly et al | Farbehi et al | Forte et al | McLellan et al | Tucker et al | Zhao et al | Li et al | Integrated dataset |
| --- | --- | --- | --- | --- | --- | --- | --- | --- |
| Publication year | 2018 | 2019 | 2020 | 2020 | 2020 | 2020 | 2020 | This paper |
| Accession number | ArrayExpress:<br>E-MTAB-6173 | ArrayExpress:<br>E-MTAB-7376 | ArrayExpress:<br>E-MTAB-7895 | ArrayExpress:<br>E-MTAB-8810 | dbGAB:<br>phs001539.v1.p1 | NCBI:<br>GSE152583 | NCBI:<br>GSE155468 |  |
| Single-cell technology | 10x Genomics | 10x Genomics | 10x Genomics | 10x Genomics | 10x Genomics | 10x Genomics | 10x Genomics |  |
| Organism | Mus musculus | Mus musculus | Mus musculus | Mus musculus | Homo sapiens | Mus musculus | Homo sapiens | Mus musculus |
| Tissue | Heart | Heart | Heart | Heart | Heart | Aorta | Aorta | Heart |
| Biological sex (age) | Female/male<br>(10 weeks) | Male<br>(8-12 weeks) | Male<br>(10-12 weeks) | Female/male<br>(10 weeks) | Male<br>(Avg. age: 50.7 years)<br>Female<br>(Avg. age: 52.5 years) | Male<br>(10 weeks old) | Male – 5<br>(Avg. age: 61.4 years)<br>Female<br>(Avg. age: 70 years) | Female/male<br>(8-12 weeks) |
| Groups | WT (*n*=4) | **Total interstitial population (TIP)**<br>Sham day 7 (*n*=4)<br>MI day 3 (*n*=4)<br>MI day 7 (*n*=4) | MI day 3: (*n*=1)<br>MI day 7: (*n*=1)<br>Sham day 7: (*n*=2) | **WT:** none<br>treatment + saline<br>treated<br>**AngII:** Angiotensin II stressed<br>(*n*=4M and 4F per group) | **LA:** left atrium (3F, 3M)<br>**LV:** left ventricle (3F, 3M)<br>**RA:** right atrium (4F, 2M)<br>**RV:** right ventricle (3F, 3M) | **Sham:**<br>Heat-inactive<br>Elastase 14 days (*n*=3)<br>Elastase-7 days (*n*=3)<br>Elastase-14 days (*n*=3) | **Control:**<br>Non-aneurysmal<br>disease control (*n*=3)<br>**ATAA:**<br>Ascending thoracic<br>aortic aneurysm (*n*=8) | **Healthy/Control :**<br>Control (McLellan et al., 2020)<br>WT (Skelly et al., 2018)<br>Sham d7 (Farbehi et al., 2019)<br>Sham d7 (Forte et al., 2020)<br>**AngII-Stressed:**<br>AngII (McLellan., 2020)<br>**MI-day3:**<br>MI-day 3 (Forte et al., 2020)<br>MI-day 3 (Farbehi et al., 2019)<br>**MI-day7:**<br>MI-day 7 (Forte et al., 2020)<br>MI-day 7 (Farbehi et al., 2019) |

The home page of the portal provides salient details of the datasets utilised and navigation to the data explorer (https://clara.baker.edu.au). The ***Datasets*** tab contains pertinent information relating to the datasets, including experimental groups and numbers, single-cell technology used, and other parameters that are important for interpretation and consideration of the data (also see Table 1). This tab also directs users to the original journal publications and allows users to download datasets from ArrayExpress or NCBI databases if required. As an educational tool, we have included a ***Cell type definitions*** tab. The glossary provides a brief description of cardiac cell types and key genes used to annotate these cell populations in the datasets. The *Data Analysis pipeline* in the ***Methods*** tab provides a detailed description of the bioinformatics workflow used to prepare scRNA-seq data within the web portal (also see Figure 1). The marker genes used to manually annotate cell populations identified in scRNA-seq datasets can be found under the *Cell Annotation* sub-section. Access to the data browser is provided by the ***Go to portal*** button in the CLARA home page and the ***Portal*** tab. This initiates an application that allows interactive data exploration. On the left panel, the dataset and gene to be displayed can be selected. The tabs on top of the Data Browser allow examination of: (i) gene expression of all cells; (ii) gene expression by condition; (iii) cell type marker genes; (iv) differential expression within cell types dependent upon context; (v) ligand-receptor signalling network for a selected ligand-receptor pair between cell types. The ***Gene expression*** tab allows the distribution of gene expression to be visualised for a chosen gene. A gene is selected by typing the gene symbol in the *gene* field, and pressing *Submit*. For example, entering *Csf1r* in ‘Mouse - Heart (McLellan *et al*., 2020)’ dataset, outputs three plots (Figure 2). The first two plots consist of points representing individual cells projected into two dimensions based on local transcriptional similarity using the tSNE algorithm; this is a method that provides a high-level visualization of cellular heterogeneity, with transcriptionally similar cells being clustered tightly in a two-dimensional space. The first plot (top left panel), is a tSNE plot where each cell population and corresponding sub-clusters are labelled and coloured. The second plot (top right panel), titled with the entered gene on top, displays a tSNE plot with the gene expression level of *Csf1r* mapped onto cells using a grey-red (low-high) colour gradient. The third plot (bottom panel), displays a violin plot – namely, a rotated kernel density plot – showing expression of *Csf1r* across each cell population. Together these three plots show that, in this example, *Csf1r* expression is primarily restricted to macrophages. These plots can be downloaded by pressing the ***Download plot*** buttons.

**Figure 1.**
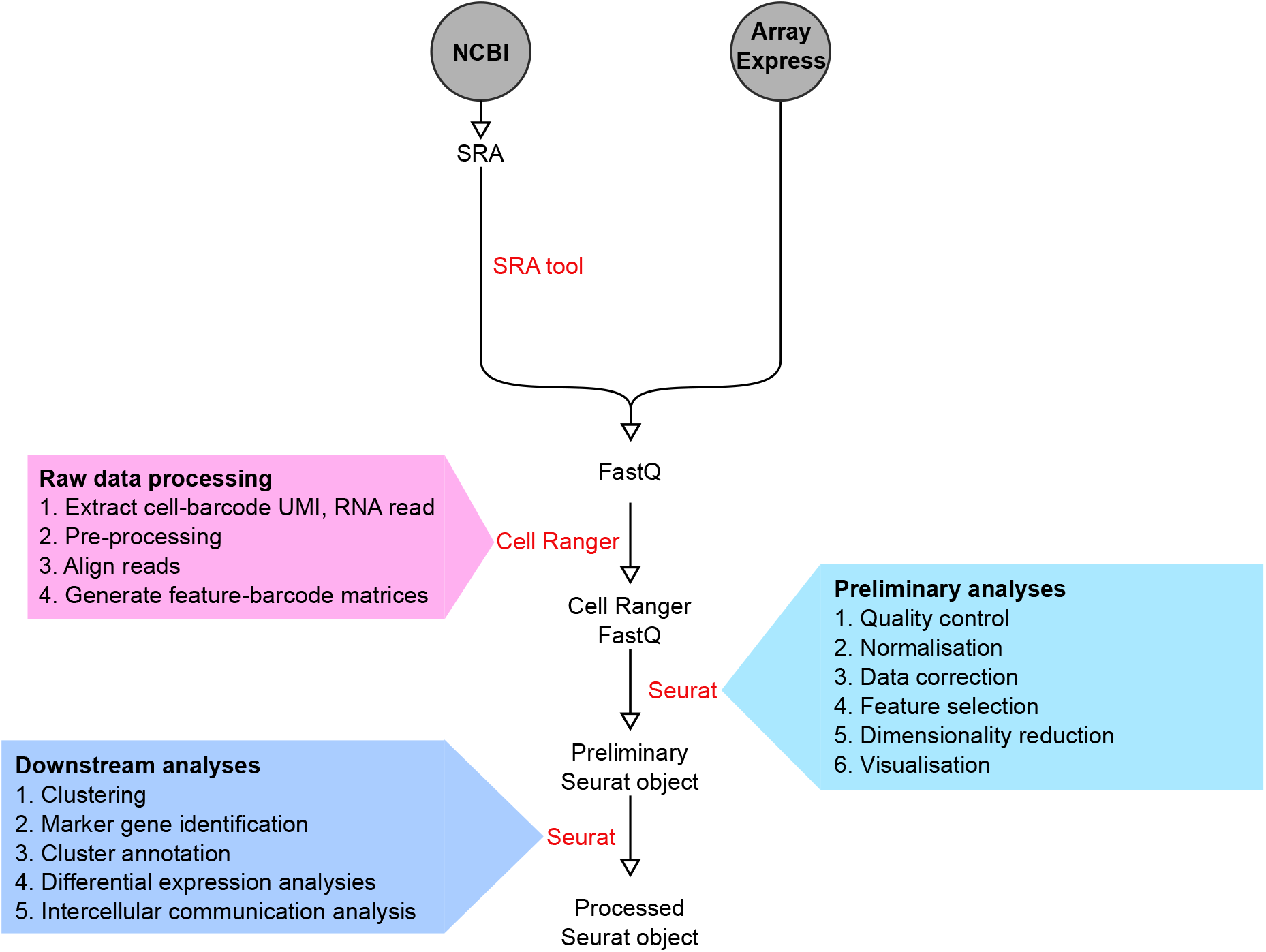
Bioinformatics workflow summarising the preparation of datasets for the portal. The raw data files were downloaded from NCBI and Array Express databases in SRA or FastQ formats, respectively. SRA files downloaded from NCBI were converted into FastQ file format using sratoolkits v.2.10.9. Subsequent FastQ files were then processed and analysed using Cell Ranger (10x Genomics) software to extract cell-barcodes, UMI and RNA reads. These RNA sequencing reads were then aligned into reference genomes to quantify transcript levels in each cell and create feature-barcode matrices. The downstream analyses scRNA-seq data were carried out in R statistical software using the Seurat R package. Figures were primarily generated using Seurat and ggplot2 R packages. See Expanded Methods in the Supplementary Materials for further details.

**Figure 2:**
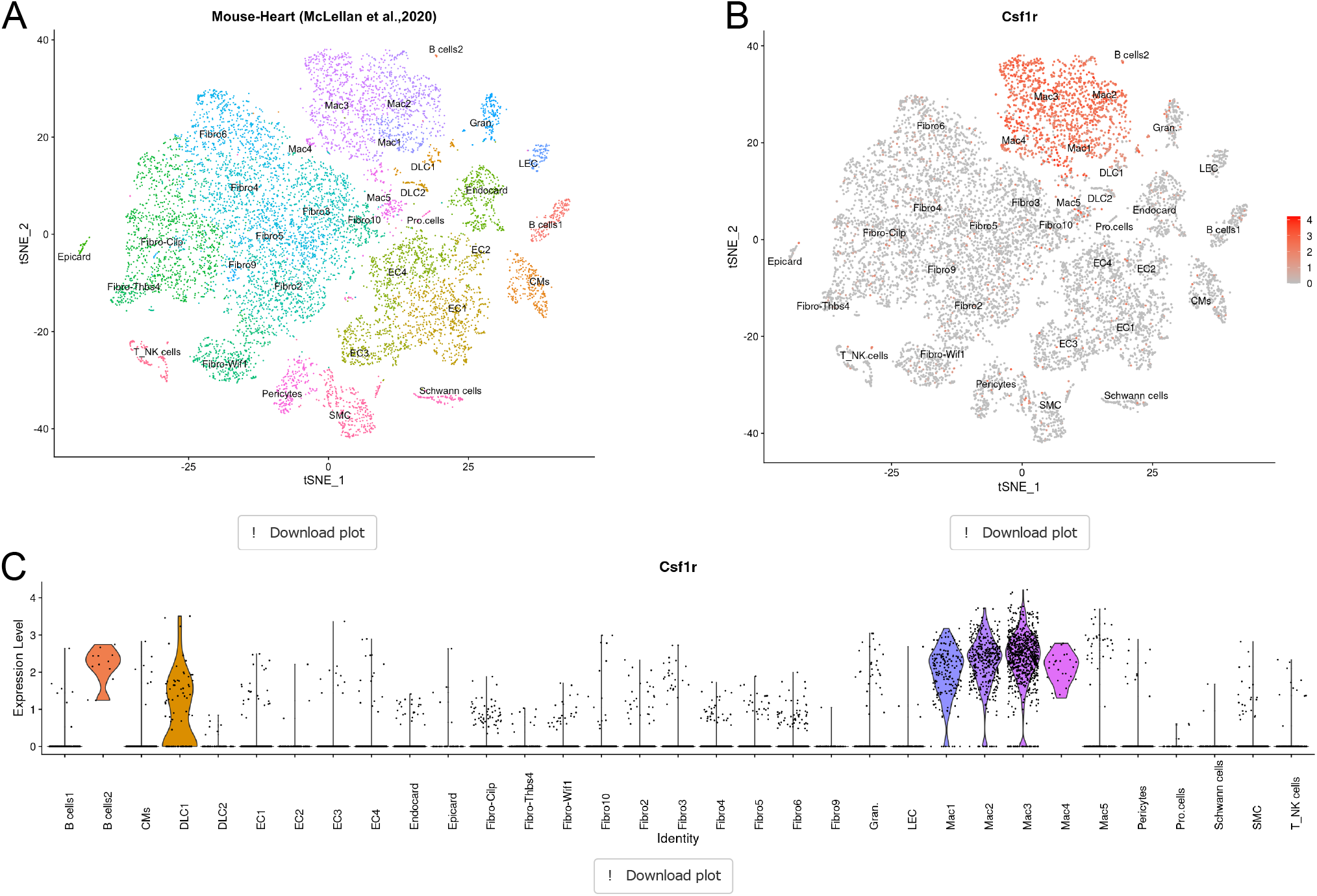
Visualization of gene expression. **(A)** tSNE projection of all cardiac cell populations and corresponding sub-clusters identified in *McLellan et al., (2020)* dataset. Using Csf1r as an example, the transcript level *Csf1r* gene is visualized in the tSNE plot with heat-map indicating relative gene expression level (red =high, grey=low) **(B)** and in Violin plots **(C)**.

The ***Gene expression by condition*** tab enables consideration of genes in context of a physiological stress. Here, the dataset and gene of choice is selected as done for the *Gene expression* tab. The data is also presented in a similar manner with tSNE projections for the control and stress condition followed by a violin plot (Figure 3). This is useful when examining genes which are up- or down-regulated following cardiac stress. For example, a query of the gene periostin (*Postn*)—a gene that is widely implicated in cardiac fibrosis— in the ‘Mouse - Heart (McLellan *et al*., 2020)’ dataset, shows that some Fibroblast-Wif1, pericytes and Schwann cells are enriched for *Postn* transcripts in control (wild type, “WT”) samples. However, in the stressed context (AngII), most cell populations are enriched for *Postn* transcript. Similar patterns are visible for the other mouse datasets where different stressors are present, demonstrating the utility of this interface for determining shifts in gene expression.

**Figure 3:**
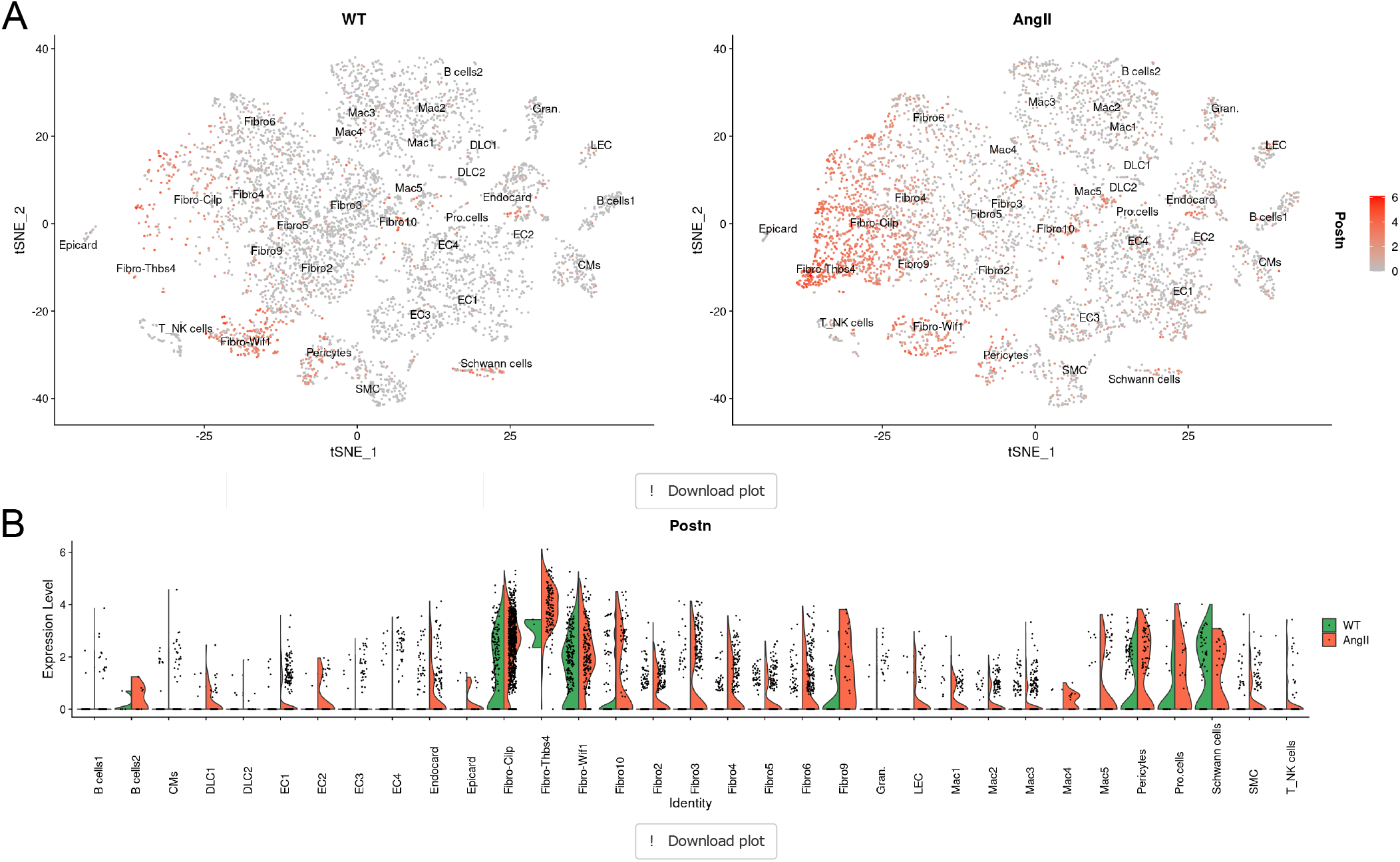
Visualization of gene expression in different contexts. **(A)** tSNE projections for the control (WT) and stressed (AngII) conditions visualizing the expression of *Postn* (periostin) gene in *McLellan et al., (2020)* dataset (red =high, grey=low) followed by **(B)** violin plots (green=control, red=stressed).

We have also included a ***Marker gene expression*** tab to examine cell type specific gene expression patterns. Here a selected cell type is displayed within a tSNE projection and a table summarises statistics relating to the top distinct genes for that cell population (Figure 4). The order of which genes are displayed can be changed by sorting column headers; however, by default, genes are ordered according to *average log fold change* (avg_logFC) relative to all other cells of the dataset.

**Figure 4:**
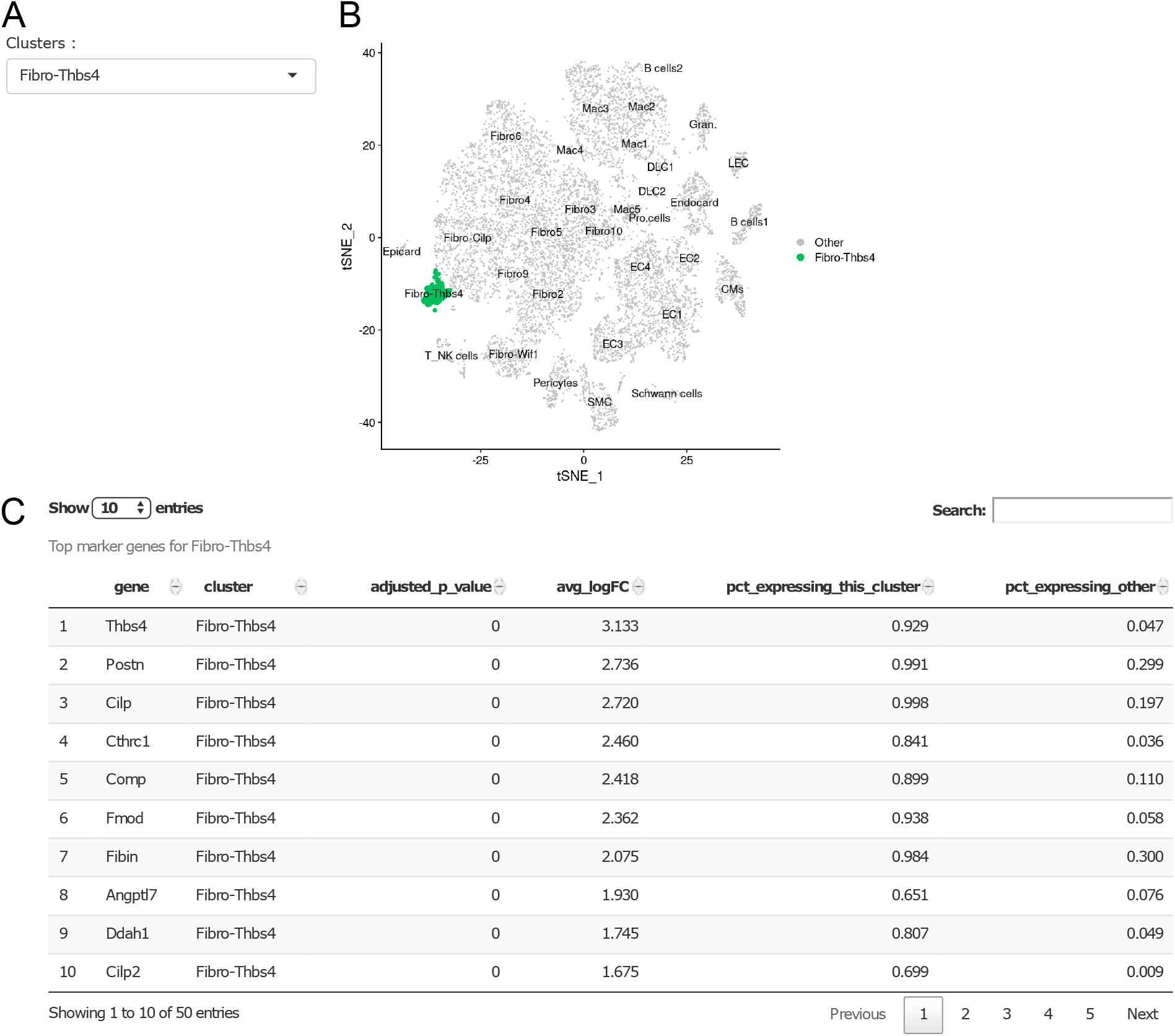
Marker genes for distinct cell populations. **(A)** drop-down menu for selecting a cell population within a dataset. **(B)** tSNE plot highlighting cells corresponding to selected ‘Fibro-Thbs4’ cell population in *McLellan et al., (2020)* dataset, used as an example here. **(C)** Table summarizing top distinct genes expressed in ‘Fibro-Thbs4’ cell population. Note: by default, genes are ordered according to average log fold change (avg_logFC) relative to all other cells in the dataset.

Context dependent gene expression can be explored by selecting the ***Differential expression*** tab. This allows selection of a cell type and a context of interest. For example, using the ‘integrated heart dataset’ (Mouse-Heart Integrated 2020) and selecting ‘pericytes 1’ and ‘Healthy-control vs AngII-stressed’ displays pericytes 1 within tSNE projections with a table summarising the top differentially expressed genes below (Figure 5). The genes within the table are ordered according to p-value (pval) by default without consideration of whether the gene is up- or down-regulated. However, the table can be reordered to determine top up-regulated genes (*Mfap4, Col1a1, and Col3a1*) and down-regulated genes (*Hspa1a, Hspa1b, and Tpi1*) by manipulating the log2 fold change (log2FC) column. Specific genes of interest within the table can also be queried by entering the gene symbol in the *Search* field.

**Figure 5.**
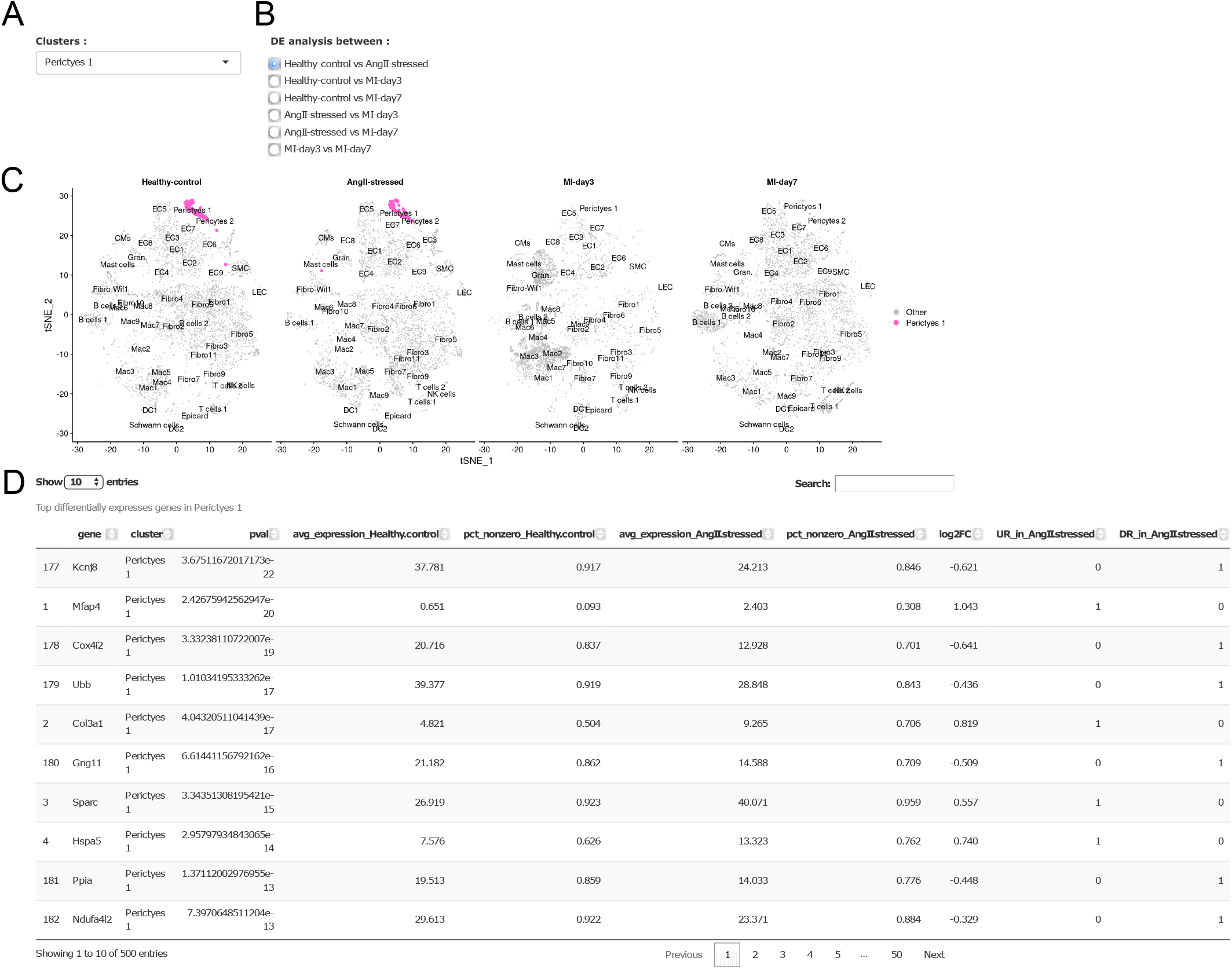
Differential gene expression between groups. **(A)** drop-down menu for selecting a cell population within the dataset. **(B)** Radio buttons for choosing conditions to compare by differential expression testing. **(C)** tSNE projections using the *Mouse-Heart Integrated 2020* dataset and ‘pericytes 1’ as an example. tSNE projections highlight ‘pericytes 1’ in Healthy-control and AngII-stressed groups. **(D)** table summarising list of top differentially expressed genes between Healthy-control and AngII-stressed groups in ‘pericytes 1’ population. Note: by default, genes are ordered according to p value (pval) without consideration to whether genes are up- or down-regulated.

Finally, putative intercellular communication networks for selected ligand-receptor pairs can be examined by selecting the ***Ligand-receptor signalling networks*** tab. Here we define ligand-receptor connections based on a dataset of human ligand-receptor pairs [9]. This tab allows selection of a ligand-receptor pair in a context of interest within a dataset. For example, selecting ligand receptor pair ‘VEGFA-KDR’ and WT in ‘Mouse - Heart (McLellan *et al*., 2020)’ dataset displays a chord plot summarizing the VEGFA-KDR communication network, as inferred from gene expression patterns, within cardiac cell populations in homeostasis (Figure 6). Arrows represent signalling direction from ligand to receptor between cardiac cell populations, with arrow thickness and colour intensity proportional to the strength of the connection. The line colour reflects the cell population producing transcript coding for the ligand. For example, arrows emerging from cardiomyocytes highlight the remarkable expression of *vascular endothelial growth factor A* (VEGFA) in cardiomyocytes which links with endothelial, endocardial, epicardial and LEC cell populations via the corresponding receptor *kinase domain insert receptor* (KDR). This signalling network clearly exhibits the significant contribution of cardiomyocytes in this micro-environment to maintain endothelial cell growth and function.

**Figure 6.**
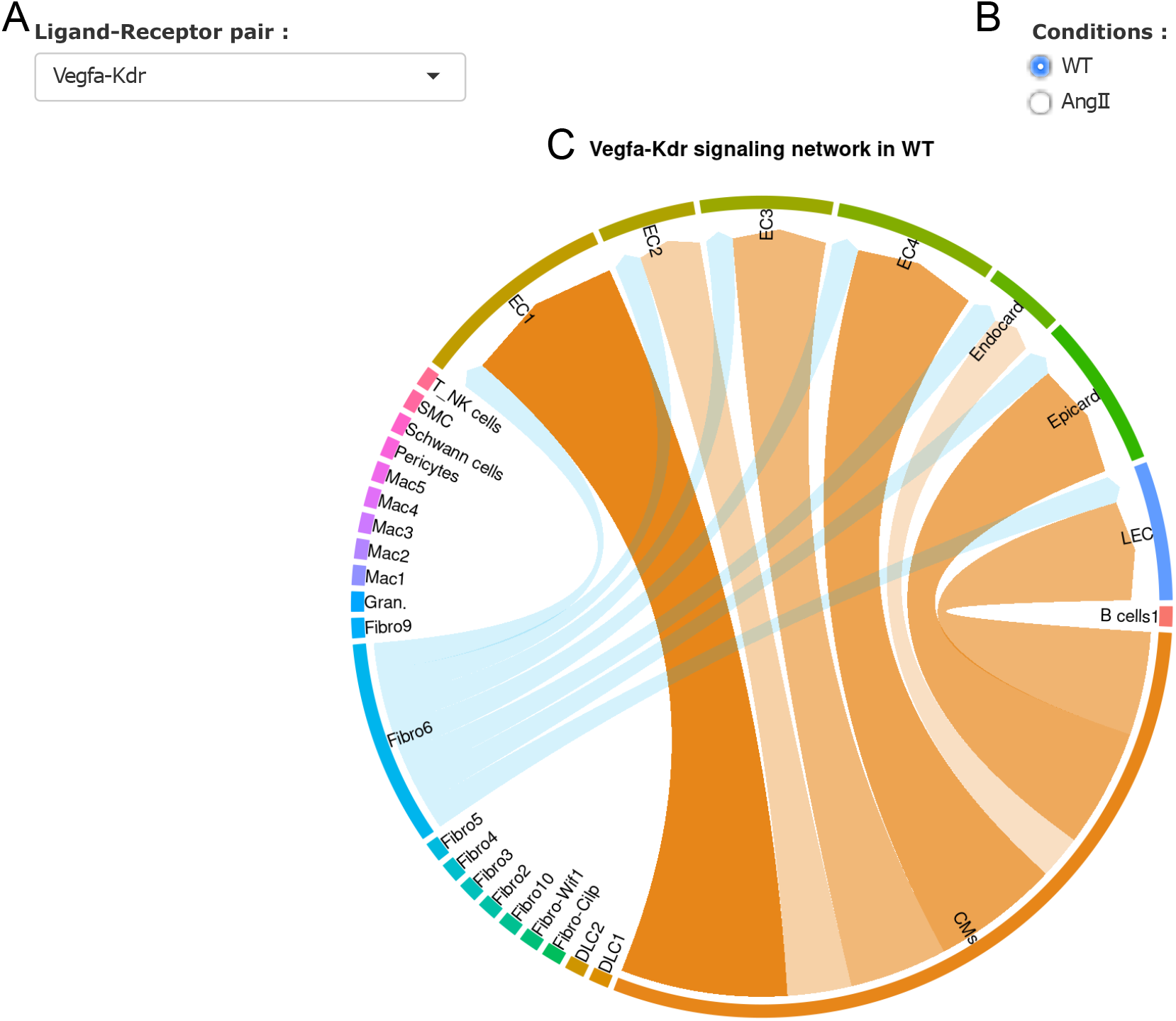
Visualization of Intercellular communication networks. **(A)** drop-down menu for selecting ligand-receptor pairs. **(B)** List of conditions within the selected dataset. **(C)** Chord plot summarising the putative signalling network for selected ligand-receptor pair. Here, we show *Vegfa-Kdr* pair in homeostasis *McLellan et al., (2020)* dataset. Arrows represent potential connections between cell populations, with arrow thickness and colour intensity proportional to the strength of the connection. The line colour reflects the cell population producing transcript coding for the ligand.

## DISCUSSION

With increasing numbers of studies incorporating single-cell omic datasets, intuitive and highly accessible tools are urgently needed for data exploration and to aide comprehension of these rich data resources by the wider research community. We present CLARA as a resource to achieve this and help further our understanding of the diverse cellular landscapes that form cardiovascular tissues. We have calibrated the presentation of CLARA to help inform individuals who are both novice and expert in the fields of cardiovascular single-cell transcriptomics, and are interested in cardiovascular cell biology in the context of health and disease. A key feature of CLARA enabling this goal is the incorporation of elements that aim to help users understand the diverse cell types within these datasets in addition to how the data was generated.

While a number of single-cell data portals are currently available, to our knowledge CLARA is the first portal focused on cardiovascular cell systems. Notable examples of alternative portals include *Single Cell Portal* (https://singlecell.broadinstitute.org) and the *Heart Cell Atlas* (https://www.heartcellatlas.org/), affiliated with the Human Cell Atlas consortium. While these portals are powerful tools, they are limited by presenting few data exploration methods (both Single Cell Portal and Human Cell Atlas) and lacking perturbations to the tissue (in the case of Heart Cell Atlas). A distinguishing feature of CLARA is the inclusion of both cardiac and aortic data from humans and mice from a number of physiological stress contexts. Datasets absent from the CLARA portal were those where raw data was unavailable at the time of developing CLARA, or where characteristics of the data precluded them from inclusion (for example, low cell numbers, low cellular heterogeneity, and/or absence of key experimental details). In addition, we have incorporated different means to visualize and explore data which incorporates features provided by other portals, such as querying cell-specific gene transcript levels, as well as novel visualization modalities for viewing intercellular communication networks. Finally, we sought to include information that may assist those interested in performing single-cell analyses independently, such as genes used for defining cell populations and overviews of data analysis pipelines.

However, a number of limitations of CLARA are worth noting. First, we down-sampled datasets presented in CLARA to ~10,000 cells for each individual datasets and 16,000 cells collectively for the integrated analysis. While this greatly enhanced performance of the portal, it may limit cellular heterogeneity represented in the dataset with potential loss of very small cell populations. Second, the datasets selected differed in animal strains, sample sizes, and protocols used to extract cells. These factors bestow challenges for comparing datasets. While we are unable to control these parameters, here we have attempted to minimize the impact of bioinformatics pipelines by reanalysing all datasets using a similar workflow and with recent releases of analysis software.

Future iterations of CLARA will have expanded capabilities and a continually updated list of included datasets. Datasets included will not be restricted to single-cell transcriptomics, and other single-cell omic data, such as *assay for transposase-accessible chromatin with sequencing* (ATAC-seq), will be incorporated. Where appropriate, we will add to existing, or create new, integrated maps of cellulomes (for instance human heart or mouse aorta) as we have done for the mouse heart in the current edition of CLARA. Second, we plan to incorporate data from other organ systems—for example the kidney and liver—which impact cardiovascular tissues. Third, we will incorporate features to examine sex-specific differences between cell types and conditions, when more studies have considered this important variable. Finally, we will update our analysis pipelines as improved approaches emerge. These include, for example, alternative databases such as *CellphoneDB* [10] for mapping ligand-receptor pairings and intercellular interactions.

In summary, CLARA enables intuitive exploration of cardiovascular cellular landscapes by visualisation of single-cell transcriptomic datasets. We anticipate that this portal will help diverse groups of individuals to interact with the rich data resources presented in CLARA to learn about cardiovascular cell populations and genetic systems. We hope to grow upon this first iteration of CLARA in the future by incorporating additional features and diverse high-dimensional datasets to further our understanding of cardiovascular cell systems in health and disease.

## METHODS

The interactive interface within CLARA (CardiovascuLAR Atlas) is implemented in R statistical software using the shiny R package [11]. The processed scRNA-seq datasets used in CLARA are stored as Seurat Objects. These objects were created using the Seurat R package [12], provide a scalable and memory efficient data format for scRNA-seq data, and integrate into R environments for visualization. CLARA contains six scRNA-seq datasets that can be easily explored on a graphical web interface for research and educational purposes. For a detailed description of datasets and analysis pipelines used in CLARA see Expanded Methods in the Supplementary Materials.

---

**Table S1: Summary table of processing parameters for each dataset.**

**Table S2: The number of cells visualized per dataset in CLARA.**

