## Supplementary Materials for "CLARA: A web portal for interactive exploration of the cardiovascular cellular landscape in health and disease"

### EXPANDED METHODS

#### scRNA-seq datasets

CLARA contains data from multiple recently published single-cell RNA-seq studies that examined cardiovascular landscapes in both human and mouse with the presence of different physiological stressors. To ensure the data diversity, we selected 4 mouse heart datasets [2–5], one human heart dataset [6], one mouse aorta dataset [8], and one human aorta dataset [7] from NCBI and Array Express databases published within 4 years' timeframe. Further, an integrated mouse heart dataset was developed by integrating selected samples from 4 mouse heart datasets [2–5] to explore a comprehensive cardiac cellular map of mouse hearts. Subsequently, in the CLARA web portal we facilitate users to explore 5 scRNA-seq datasets from individual studies [3, 4, 6–8] and one dataset from our integrated analysis (Mouse-Heart Integrated 2020). Each dataset revealed cellular and transcriptional diversity in different conditions of the heart and aorta. See Table 1 and https://clara.baker.edu.au/datasets/ for more details about single-cell RNA-seq datasets used in CLARA. This includes publication details, single-cell technology used, tissue context, experimental groups and number of samples. For mouse or human heart, Farbehi *et al*., (2019) dataset explores the cell interaction in murine hearts at days 3 and 7 of myocardial infarction (MI) surgery and day 7 post-sham. McLellan *et al.,* (2020) uncovered murine cardiac cellular ecosystem in Angiotensin II treatment (AngII). Tucker *et al.,* (2020) provided extensive cellular landscape of the 4 chambers of the non-failing human heart. Regarding human or mouse aorta datasets, Li *et al*., (2020) characterized the cellular network in the ascending thoracic aorta after aneurysm (ATAA). The cellular response and heterogeneity of mouse aorta during abdominal aortic aneurysm (AAA) progression was outlined in Zhao *et al*., (2020) by comparing infra-renal abdominal aorta (IAAs) at days 7 and 14 after elastase exposure (Elastase 7 days and Elastase 14 days) with the Sham group. In terms of the integrated mouse dataset, we defined four conditions based on initial conditions in individual datasets. Healthy-control condition included the control group from Skelly *et al*., (2018), day 7 sham from Farbehi *et al*., (2019) and Forte *et al*., (2020), and non-treated and saline-treated samples from McLellan *et al*., (2020). AngII group from McLellan *et al.,* (2020) was defined as AngII-stressed. MI-day3 and MI-day7 contained corresponding samples from Forte *et al*., (2019) and Farbehi *et al*., (2020).

#### scRNA-seq data pre-processing

The corresponding raw data files for selected scRNA-seq datasets were retrieved from NCBI or Array Express databases. The SRA files downloaded from NCBI were first converted into FastQ file format using SRA toolkit v2.10.9. Then, Cell Ranger versions 3.1.0, 4.0.0 and 5.0.0 (10X Genomics) were used to process raw sequencing data before subsequent analyses. This pipeline used FastQ files to extract cell-barcodes, UMI and RNA reads. These RNA sequencing reads were then aligned into the mm10/GRCm38 transcriptome for mouse and GRCh38 transcriptome for human datasets, quantifying the expression of transcripts in each cell to create feature-barcode matrices. Datasets that were created using single-nuclei experiments were aligned to a pre-mRNA reference genome downloaded from the Cell Ranger software to account for pre-mature RNA. Single-nuclei transcriptomes usually contain a higher fraction of reads mapping to intron regions than single-cell transcriptomes. The analyses of processed scRNA-seq data were carried out in R version 3.6 and 4.0.1 using the Seurat suite versions 3.2.0 and 4.0.1.

For each dataset, initial quality control filtering metrics were applied. Cells with fewer than a given number of detected expressed genes and genes that were expressed in less than a given number of cells were first filtered out. Further, cells with over a given percentage of raw UMIs mapping to mitochondrial genes were filtered out to control for dying or damaged cells. The pre-processing parameters for each dataset are detailed in the Supplementary materials (See Table S1).

**Clustering and marker gene identification**

UMIs were normalized to counts-per-ten-thousand, log-transformed, and a set of highly variable genes was identified by gating for mean expression level and dispersion level in a per-data-set manner. The log-normalized data were then scaled, with variation due to the total number of UMIs regressed out using a linear model. Principal component analysis was run on the scaled data for the set of previously defined highly variable genes per dataset. In order to identify the number of principal components (PCs) to use for clustering, we used elbow plots implemented in the Seurat R package [12].

The steps and parameters used for dimensionality reduction and clustering for each dataset are available in Supplementary Materials Table S1. The principal component analysis was used to determine the optimal number of principal components for each dataset which is later used for non-linear dimension reduction such as t-SNE. The optimal resolution for each dataset was determined to avoid over-clustering and to be consistent with the original publication. The resolution parameter for clustering, which determined the number of returned clusters, was decided on a per-data-set basis after considering clustering output from a range of resolutions. The cells and clusters were then visualized on a t-SNE dimensionality reduction plot generated using the optimal number of PCs and resolution.

Moreover, statistical methodologies implemented in Seurat R package were used to control potential batch effects present in some datasets and to integrate multiple datasets together. For example, the canonical correlation analysis (CCA) was performed on Li *et al*., (2020) dataset to remove unwanted variability between samples. Also, Seurat V3 integration (standard workflow) [13] was used to integrate multiple single-cell datasets to create the Mouse-Heart integrated 2020 dataset.

After cell clustering, marker genes per cell cluster were identified using 'FindAllMarkers' Seurat R function per dataset. Identified cell clusters were then annotated based on known marker genes for consistency with original publications. Also see cell annotation methodology and marker genes used in CLARA at https://clara.baker.edu.au/methods/cell-annotation/.

**Down-sampling scRNA-seq datasets**

After pre-processing, the number of cells per dataset was between 5,000-60,000 with genes per dataset being ~16,000-28,000 (see supplementary Table S1 for more details). For better visualization and memory handling of the CLARA shiny application, we down-sampled each single-cell RNA-seq dataset and randomly selected 10,000-16,000 cells per dataset to create smaller Seurat objects. The number of cells visualized in CLARA per dataset is detailed in Supplementary Table S2.

**Differential expression analysis**

In order to identify differentially expressed (DE) genes in each dataset, we first filtered genes expressed in at least 10% of cells in at least one of the groups being compared. This was performed for each cell type separately. The differential expression analysis method MAST [14] was used to identify DE genes between groups considering cellular detection rate as a covariate, which performed well in the single-cell RNA-seq differential expression benchmarking study conducted by Soneson and Robinson [15].

**Intercellular communication analysis**

In order to represent potential intercellular communication between cardiac cell populations based on a given ligand-receptor gene pair, we used human ligand-receptor pairs published in Ramilowski *et al*., (2015). The BioMart database version Ensembl 101 was used to obtain mouse orthologs of those human ligand-receptor pairs using the biomaRt R package. A ligand/receptor with non-zero expression in more than 20% of cells in a particular cell population was considered an "expressed" ligand/receptor.

To construct the putative cell-cell signalling network for a given ligand-receptor pair in a given condition, the average expression per cell population was first calculated. Cell populations with less than 5 cells were discarded from this analysis. In the ligand-receptor signalling network visualization, we linked expressed ligand with its corresponding receptor between and within cell populations. The signalling direction from ligand to receptor is indicated by an arrow with thickness and colour intensity proportional to the strength of the potential interactions between cell populations. The potential strength of a ligand-receptor interaction was calculated based on the expression values of both ligand and receptor genes in corresponding cell populations. The line colour reflects the cell population producing transcript coding for the ligand.

**Table S1: Summary table of processing parameters for each dataset.**

| Dataset | Cell Ranger version | Reference genome | R/Seurat versions | Seurat object initialisation | QC parameters | Number of cells and genes after QC | Normalisation Method and Integrations | Number of PCs and clustering resolution |
|---|---|---|---|---|---|---|---|---|
| McLellan et al., 2020 | 4.0.0 | mm10-2020-A | R 3.6; Seurat 3.2.0 | min.cell=3 min.features=200 | nFeature_RNA > 200<br>percent.mt < 20 | Cells - 37195<br>Genes- 18857 | **Normalisation:** LogNormalise | PCs - 40<br>Res - 1.2 |
| Farbehi et al., 2019 | 4.0.0 | mm10-2020-A | R 3.6; Seurat 3.2.0 | | nFeature_RNA > 200<br>percent.mt < 10 | Cells- 22,178<br>Genes - 19,097 | **Normalisation:** LogNormalise | PCs - 40<br>Res - 1.2 |
| Integrated 2020 | 4.0.0 | mm10-2020-A | R 3.6; Seurat 3.2.0 | | nFeature_RNA > 200<br>nFeature_RNA < 6000<br>percent.mt < 20 | Cells - 127,348<br>Genes - 20,801 | **Normalisation:** LogNormalise<br>**Integration:** CCA | PCs - 50<br>Res - 1.2 |
| Tucker et al., 2020 | 3.1.0 | GRCh38-2020-A_premrna | R 4.0.1; Seurat 4.0.1 | | nFeature_RNA > 100<br>nFeature_RNA < 10000<br>nCount_RNA< 25000<br>percent.mt < 5 | Cells - 251,004<br>Genes - 30,057 | **Normalisation:** SCTransform<br>**Integration:** reference based + RPCA | PCs - 50<br>Res - 0.5 |
| Zhao et al., 2020 | 3.0 | mm10, version 1.2.0 | R 4.0.1; Seurat 4.0.1 | | nFeature_RNA > 200<br>nFeature_RNA < 7000<br>percent.mt < 10 | Cells - 4,854<br>Genes- 16,224 | **Normalisation:** LogNormalise | PCs - 40<br>Res - 0.8 |
| Li et al., 2020 | 5.0.1 | GRCh38-2020-A | R 4.0.1; Seurat 4.0.1 | | nFeature_RNA > 200<br>nFeature_RNA < 9000<br>percent.mt < 30 | Cells - 59,562<br>Genes- 27,261 | **Normalisation:** LogNormalise<br>**Integration:** CCA | PCs - 30<br>Res - 0.8 |

**Table S2: The number of cells visualized per dataset in CLARA.**

| Dataset | Total number of cells | Number of cells visualised in CLARA - selected by random sampling |
|---|---|---|
| McLellan et al., 2020 | 37,195 | 10,000 |
| Farbehi et al., 2019 | 22,178 | 10,000 |
| Integrated dataset 2020 | 127,348 | 16,000 (4000 cells x 4 groups) |
| Tucker et al., 2020 | 251,004 | 12,000 (3000 cells x 3 groups) |
| Zhao et al., 2020 | 4,854 | 4,854 (full dataset) |
| Li et al., 2020 | 59,562 | 10,000 (5000 cells x 2 groups) |
